# SARS-CoV-2 Infection Causes Strong mTORC1 Inhibition and Massive Polysome Collapse

**DOI:** 10.1101/2021.05.08.443207

**Authors:** Haripriya Parthasarathy, Divya Gupta, Dixit Tandel, Vishal Sah, Abdul Hamid Siddiqui, Abhirami P Suresh, Krishnan Harinivas Harshan

## Abstract

Viruses employ distinct strategies to ensure efficient translation of their mRNAs over the host transcripts. SARS-CoV-2 targets host mRNAs and ribosomes to favor its own protein synthesis. However, the modulation of the key signal pathways that control the host protein translation machinery during SARS-CoV-2 infection has not been sufficiently addressed. Here, by employing an early variant and a Delta variant isolate that evolved later in the pandemic demonstrates that SARS-CoV-2 infection results in massive polysome collapse starting from 24 hpi, a hallmark of global translation inhibition. Unexpectedly, eIF2α phosphorylation, commonly targeted by viruses to induce translation arrest, was not involved in the translation arrest, suggesting that SARS-CoV-2 countermeasures by the virus to suppress ISR. eIF4E phosphorylation remained unaltered during the infection, ruling out its involvement in the preferential translation of SARS-CoV-2 transcripts. We find that SARS-CoV-2 infection consistently causes mTORC1 inhibition in a comparable manner across both variants indicating that the virus likely targets mTORC1 pathway to suppress host translation. Interestingly, mTORC1 inhibition by SARS-CoV-2 did not impact the polysomal loading of ribosomal protein transcripts rpS3 and rpL26, suggesting that 5’TOP mRNAs are spared from the translation suppression and that ribosomal protein synthesis remains active during the infection. Pharmacological activation of mTORC1 did not significantly impact viral replication, suggesting that mTORC1 inhibition might be selectively restricting the host mRNAs from accessing the translation machinery, facilitating a more robust translation of viral transcripts. This study provides new insights into the molecular interactions by which SARS-CoV-2 variants, despite their different clinical outcomes, converge on a conserved mechanism to manipulate host translation regulatory pathways.

## INTRODUCTION

Severe acute respiratory syndrome coronavirus 2 (SARS-CoV-2) has emerged as the deadliest of all known coronaviruses up to date, causing more than 7 million deaths worldwide(1,2). Inside a host cell, the 5’-capped and polyadenylated viral genomic RNA undergoes multiple rounds of translation to generate two long polypeptides that are cleaved by proteases to generate non-structural proteins that assemble to form the replicase complex. Translation of sub-genomic mRNAs generate structural proteins that are temporally regulated, indicating the importance of tightly regulated translation for successful infection by SARS-CoV-2(3). Given that translation control is central to viral life cycle, perturbation of host translation emerged as a common strategy observed in SARS-CoV-2(4)

Viruses establish distinct relationships with the host translation machinery, ranging from subverting it to coaxing, or partly manipulating it. Poliovirus completely shuts down host translation to utilize the now available machinery for cap-independent translation of viral transcripts (5). Several other viruses inhibit host translation to varying degrees while allowing a set of mRNAs to translate (6,7). Yet, some other viruses such as hepatitis C virus (HCV) do not cause an apparent suppression of host translation, but still use a cap-independent mechanism for their translation. Coronaviruses that gained significance in the recent past such as SARS-CoV and MERS-CoV are known to inhibit host translation through a variety of mechanisms such as inhibition of 40S subunit and 80S formation, mRNA degradation, and disrupting protein trafficking (7–12) Extensive studies have established a potent involvement of several SARS-CoV-2 non-structural proteins in the arrest of host translation. Most notably, Nsp1 interferes with host translation through its interaction with 40S ribosomes(7,13–15). Reports also indicate SARS-CoV-2 mediated preferential destruction of host mRNAs lead to their reduced translation events (7,16). While a number of different mechanisms by which SARS-CoV-2 interferes with host translation activities have been proposed, newer reports suggest significant differences between the variants that have emerged over time (17,18)

Global translation activities in higher eukaryotes are regulated by three major pathways. Mechanistic target of Rapamycin (mTOR) pathway is the most studied of these and is known to regulate translation of a sub-set of mRNAs with a 5’ terminal oligo pyrimidine (5’ TOP) stretch (19–21). mTOR complex 1 (mTORC1) is active in metabolically active cells and promotes translation by facilitating the free availability of the cap-binding initiation factor, eIF4E (22). One of the substrates of mTORC1, eIF4E binding protein 1 (4EBP1), inhibits translation activities by sequestering eIF4E (23). mTORC1 mediated phosphorylation of 4EBP1 lowers its affinity towards eIF4E thereby making it available for cap-binding. mTORC1 also facilitates translation by phosphorylating the small ribosomal protein rpS6(24), eIF4B and helicase eIF4A through p70S6K (25). Several viruses are reported to target mTORC1 in order to suppress host translation activities(11). Inhibition of mTORC1 is known to cause a major drop in active polysomes and translation activities (20,21).

MAPKs p38 and ERK1/2 are known to regulate the phosphorylation of eIF4E through their substrate Mnk1/2(26,27). Even though eIF4E phosphorylation does not alter its affinity for the 5’ cap of the mRNAs, phosphorylated eIF4E is commonly detected in several cancers leading several researchers to hypothesize that this phosphorylation results in preferential translation of a set of mRNAs (28,29). A third mechanism of regulation of global translation is the phosphorylation of eIF2α at S52, a key event leading to reduced recycling rate of eIF2 ternary complexes that is critical for new events of translation initiations (30). The four kinases GCN2, HRI, PKR and PERK, known as integrated stress response (ISR) kinases, coordinate this phosphorylation relaying various upstream signals (31) PKR, phosphorylates eIF2α after the detection of dsRNA replication intermediates in the cytosol during viral infections. This results in severe translational suppression in the virus infected cells as demonstrated in several cases (32,33).

While the majority of studies aimed at understanding the impact of SARS-CoV-2 infection on host translation have demonstrated a major translation inhibition by the virus, our study focused on delineating the impact of the virus on host signaling pathways that have a major impact on translation. We analysed this by employing two distinct SARS-CoV-2 variants with one of the representing an early lineage isolated during the early months of the pandemic, while the other one representing Delta variant that emerged during the later phase in the pandemic. Our findings demonstrate that both variants trigger a comparable and sustained polysome dissociation beginning at 24 hrs post infection and persisting upto 96 hrs. Further assessment of pathways regulating host translation revealed that p38 MAPK was phosphorylated by both variants throughout the course of infection and its inhibition also resulted in lower viral replication. Interestingly, despite activated p38 MAPK, eIF4E phosphorylation was limited. However, both variants consistently, and strongly inhibited the mTORC1 pathway. Nevertheless, the translation of 5’ TOP transcripts continued unhindered in cells infected with both variants, indicating escape of these mRNAs from virus-mediated mTORC1 inhibition as well as translation arrest. Collectively, our studies demonstrate that SARS-CoV-2 infection causes severe arrest of host translation machinery, and this is potentially supported by a strong inhibition of mTORC1 in a selective manner to spare the key host transcripts. Importantly, these effects were conserved between the two variants despite their distinct stage of emergence and different clinical manifestations, indicating that SARS-CoV-2 employs a shared molecular strategy to modulate host translation.

## RESULTS

### SARS-CoV-2 causes major polysome collapse

Previous reports have characterized a strong suppression of translation of host transcripts upon SARS-CoV-2 infection through Nsp1-mediated host mRNA degradation. However, few have explored the influence of the viral infection on host translation apparatus and signal pathways that maintain it. We studied the effect of SARS-CoV-2 infection on polysome formation as an indicator of translation activity. We used an earlier variant isolate B.1.1.8 and a later variant isolate of Delta in this study to infect Caco2 cells(34). Both variants established robust infection as evident from nucleocapsid (N) expression (Figures 1A and D). A significant collapse of actively translating polysome pool was observed in B.1.1.8 infection as early as 24 hours post infection (hpi), (Figure 1B and C), indicating a rapid suppression of translation activities. Quantification of the polysome area demonstrated a polysome depletion by about 40% in the virus infected samples by 24 hpi against the uninfected control cells, indicating that the virus causes a marked reduction in translation activity. Notably, this reduction in the translating pool of polysomes was maintained throughout the infection course. A moderate increase in the 80S peak was observed during the infection, suggesting that the fresh rounds of translation initiation were not impacted. Delta infection also showed a comparable pattern of polysome dissociation (Figures 1E and F). Curiously, no proportionate increase in the size of the mRNPs fraction was noticed, suggesting that the loss in the polysomes did not result in large-scale degradation of the mRNA associated with them (Supplementary Figures S1A and B). Under these conditions, N protein maintained its levels, indicating that the viral translation is unaffected by the loss in polysome fractions. The possibility that the polysome collapse is an outcome of cell death due to viral infection was ruled out by the cytotoxicity assay (Figures 1G for B.1.1.8 and H for Delta).

**Figure 1:**
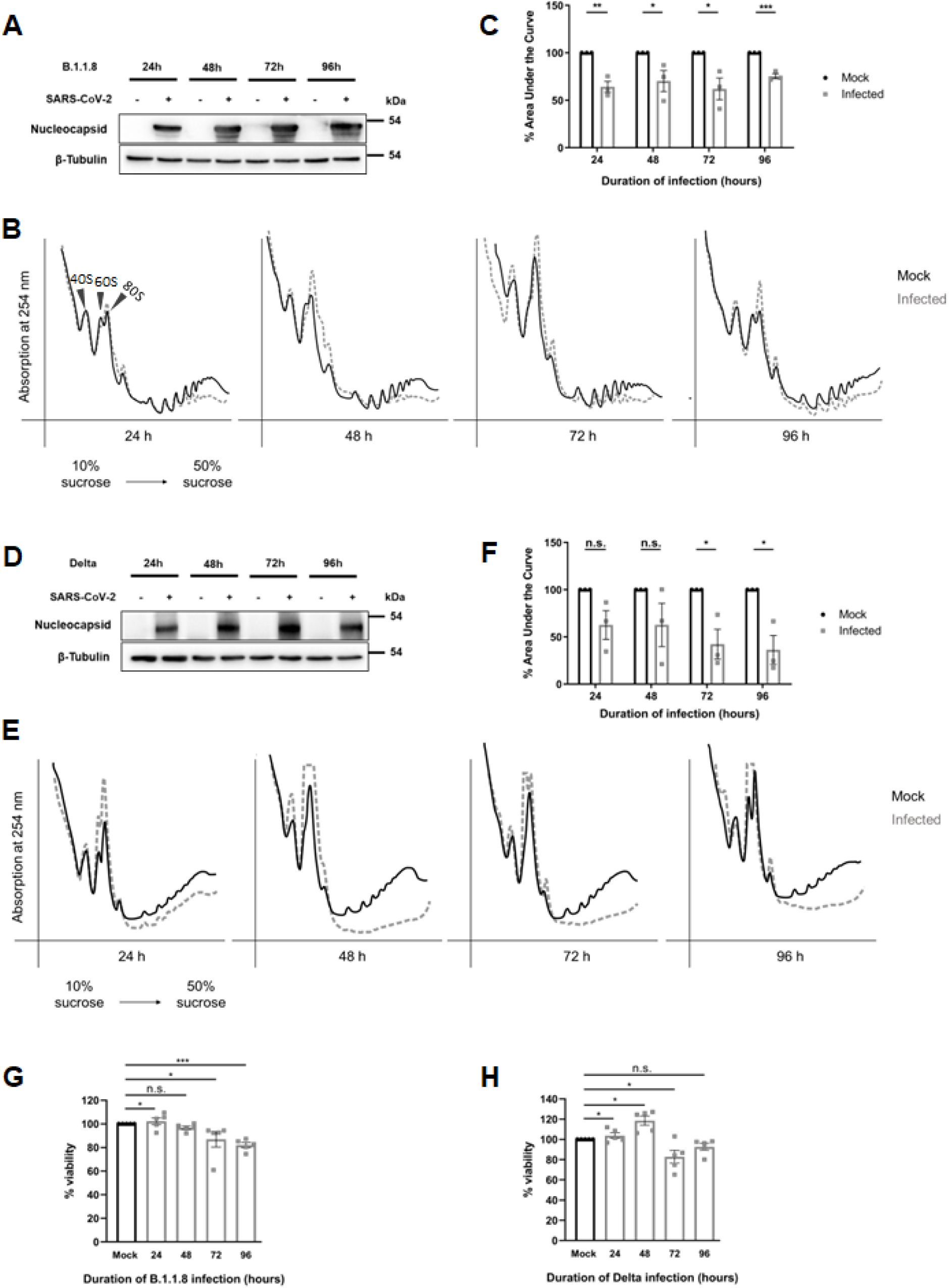
SARS-CoV-2 causes major polysomal collapse. **(A, D)** Immunoblot analysis of mock and B.1.1.8 **(A)** or Delta **(D)** infected Caco2 cells over 24, 48, 72, and 96 hrs of infection. Cell lysates were separated by SDS-PAGE and probed for SARS-CoV-2 nucleocapsid protein. **(B, E)** Polysome profiles of Caco2 cells infected with B.1.1.8 and Delta for 24, 48, 72, and 96 hrs. The cells were treated with 100 µg/mL CHX before harvesting and lysed in polysome lysis buffer. Equal quantities of lysates were layered onto continuous sucrose gradients ranging from 10-50%, subjected to ultracentrifugation, and fractionated along with measuring absorbance at 254 nm. The digitized profiles for infected and uninfected samples for each time point were overlaid to assess any change in global translation levels. The profiles were digitalized using Inkscape software. Arrows indicate inferred positions of 40S, 60S, and 80S peaks, based on known sedimentation standards (George et al., J Biol Chem, 2012) **(C, F)** Graphs showing % area under the curve (AUC) of mock and SARS-CoV-2 infected polysome profiles over 24, 48, 72, and 96 hrs of infection for B.1.1.8 and Delta respectively. The area under the curve was calculated using ImageJ software. The polysome area, excluding mRNPs fractions, was normalized to the profile’s total area. Graphs represent data from three independent sets for B.1.1.8 **(B)** and Delta **(E)**, respectively. **(G)** and **(H)** Graphs showing percentage cell viability of infected cells compared to mock at different time intervals from B.1.1.8 and Delta infection respectively. All values are plotted as mean ± SEM. p-values were calculated using Student’s t-test and represented as * and **, indicating p-values ≤ 0.05 and 0.005, respectively.

To further characterize the molecular means to this polysome loss, we studied mTOR, MAPK, and eIF2α pathways upon infection. Strong signals of N protein and viral RNA were detected by 12 hpi in both B.1.1.8 and Delta infections (Supplementary Figures S2 A-C; S3 A-C, respectively). Interestingly, none of the three pathways indicated any signs of regulation suggesting that SARS-CoV-2 infection does not result in an early inhibition of protein translation initiation (Supplementary Figures S2 D-L; Figures S3 D-L, respectively). We subsequently extended our studies during 24-48 hpi by when the virus had successfully established infection.

### SARS-CoV-2 does not induce ISR

eIF2α phosphorylation-mediated translational arrest is often used by viruses to achieve preferential translation of viral proteins (32,33). We first asked if SARS-CoV-2 infection potentially leads to eIF2α phosphorylation resulting in polysome dissociation and translational arrest. Unexpectedly, eIF2α phosphorylation remained unchanged at 24 hpi, followed by a sharp decrease from 48 hpi onwards in B.1.1.8 infection, Although much delayed, Delta infection also caused a severe eIF2α dephosphorylation. Collectively, these findings suggest that SARS-CoV-2 mediated translational suppression does not primarily involve engagement of the eIF2α pathway (Figures 2A and C).

**Figure 2:**
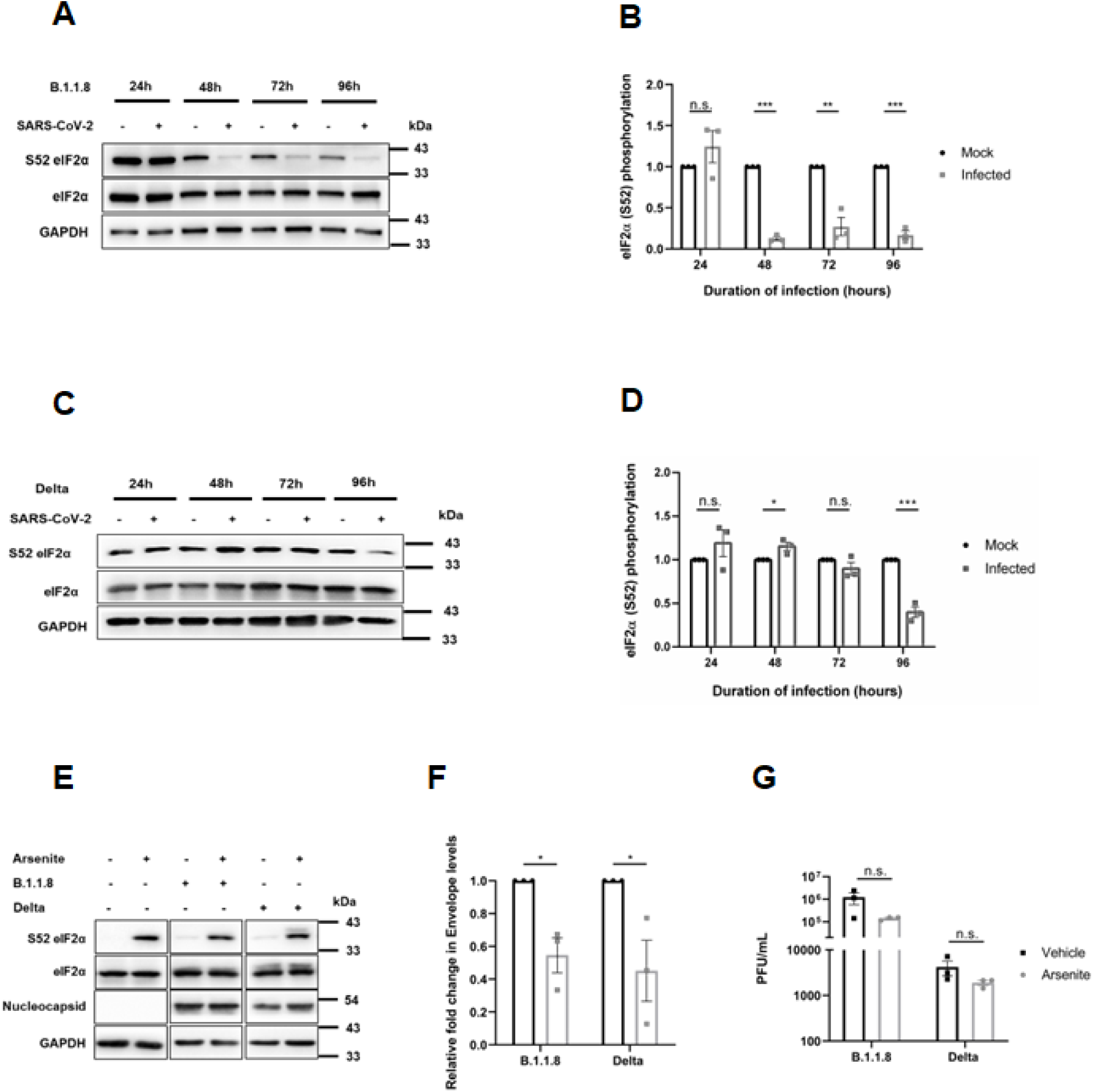
SARS-CoV-2 does not induce ISR-eIF2α axis. **(A, C)** Immunoblots showing changes in phosphorylation of eIF2α at S52, and its expression in Caco2 cells over different time points of infection with B.1.1.8 **(A)** and Delta **(C)**, respectively. **(B, D)** Densitometric analysis from the panel for B.1.1.8 **(B)** and Delta **(D)**, respectively. **(E)** SARS-CoV-2 variants infected Caco2 cells were treated with sodium arsenite at 10 μM concentration for 8 hrs starting at 16 hpi. The effect of treatment was scored by the phosphorylation status of eIF2α at S52, and the abundance of viral protein under arsenite treatment was also studied. **(F)** Relative fold change in SARS-CoV-2 viral Envelope levels in infected cells treated with either vehicle or 10 μM arsenite. Value from vehicle-treated cells were used for normalization, and the relative fold change is represented graphically. **(G)** Graph showing changes in viral titers with vehicle or arsenite treatment quantified as PFU/mL. Graphs represent data from at least three sets and are plotted as mean ± SEM. p-values are represented as *, ** and ***, indicating p-values ≤ 0.05, 0.005 and 0.0005, respectively, n.s. represents non-significant.

Inhibition of eIF2α phosphorylation by SARS-CoV-2 is an indication that the virus prefers to suppress ISR or does not want to limit the availability of ternary complexes for the interest of its own translation. Next, we asked if activating ISR would be deleterious to the virus. Caco2 cells infected with SARS-CoV-2 were treated with sodium arsenite, a general inducer of eIF2α phosphorylation and its impact on the viral titer was studied. Sodium arsenite induced strong eIF2α phosphorylation in the infected cells, but failed to reduce the levels of viral N protein (Figure 2E). However, a moderate drop in viral RNA as well as infectious viral titers were observed in the supernatant of arsenite treated cells as compared with the control (Figures 2F and G), suggesting that the viral translation is partially impacted by elevated eIF2α phosphorylation, and the virus may have evolved ways to translate under conditions of depleted canonical ternary complex.

### MAPK-eIF4E axis does not mediate polysome collapse during SARS-CoV-2 infection

Since eIF2α did not mediate polysome collapse in SARS-CoV-2 infection, we asked if the two MAPKs, ERK1/2 or p38 MAPK mediate this outcome. We first investigated the phosphorylation of ERK/2 in SARS-CoV-2 infected cells. Interestingly, as in the case of eIF2α, ERK1/2 phosphorylation remained unchanged until 24 hpi, but started to decrease from 48 hpi onwards (Figures 3 A-B and D-E). However, eIF4E phosphorylation remained unchanged throughout the infection cycle (Figures 3C and F). These results, consistent for both the variant infections, indicated that SARS-CoV-2-induced polysome collapse is not mediated through ERK1/2-eIF4E.

**Figure 3:**
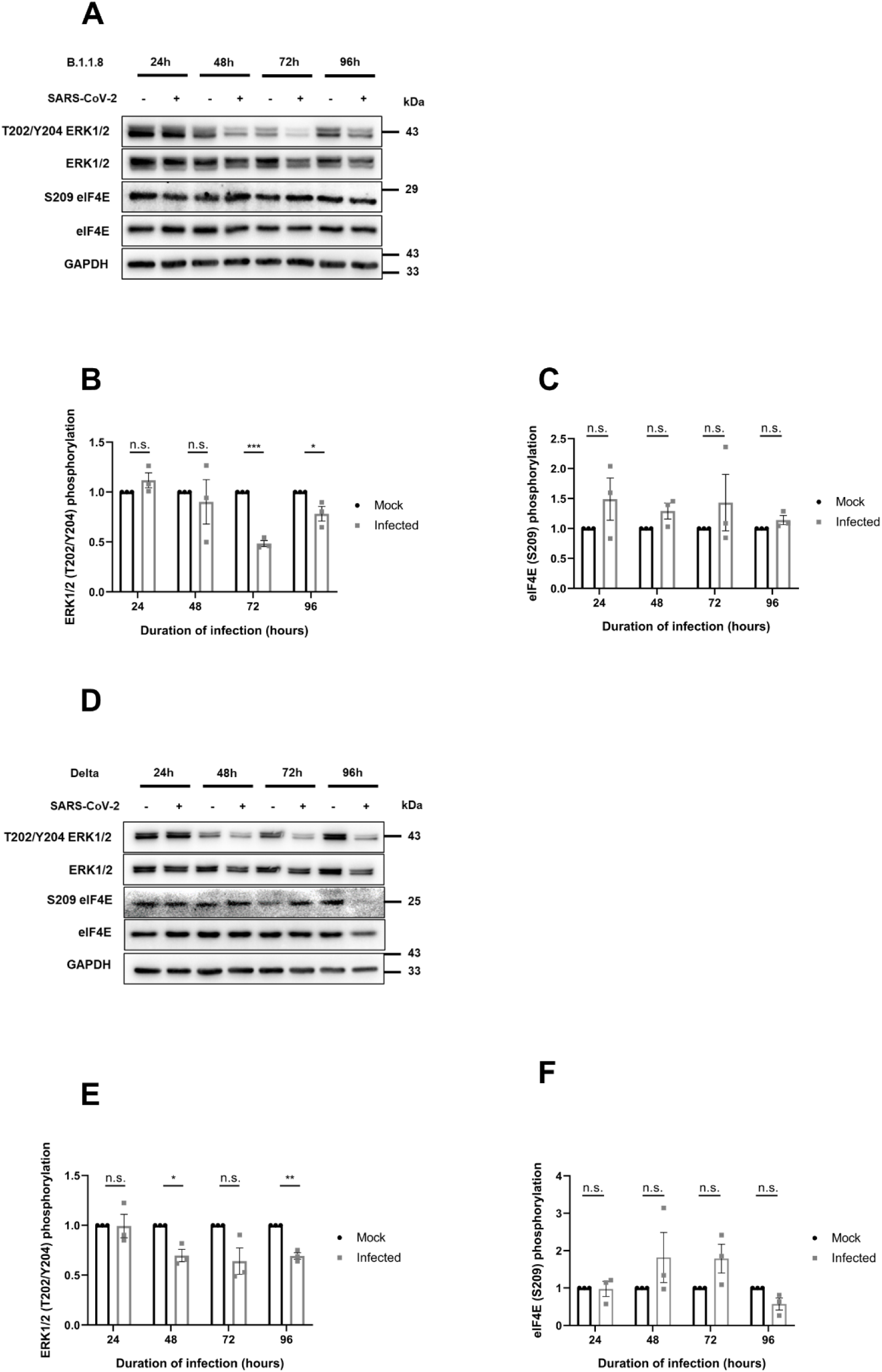
MAPK-eIF4E axis does not mediate polysome collapse during 5vfr34bCoV-2 infection. **(A)** Immunoblots depicting phosphorylation and expression status of ERK1/2 and eIF4E from Caco2 cells infected with B.1.1.8 for 24, 48, 72, and 96 hrs along with mock-infected controls. **(B)** Densitometric quantitation of T202/Y204 ERK1/2 phosphorylation from the panel in A. **(C)** Densitometric quantitation of S209 eIF4E phosphorylation from the panel in A. **(D)** Immunoblot showing changes in phosphorylation and expression status of ERK1/2 and eIF4E with Delta infection. **(E)** Densitometric quantitation of T202/Y204 ERK1/2 phosphorylation from the panel in D. **(F)** Densitometric quantitation of S209 eIF4E phosphorylation from the panel in D.

Next, we analyzed the status of p38 MAPK during the infection. Unlike ERK1/2, p38 MAPK phosphorylation was induced during SARS-CoV-2 infection from 48 hpi in both variant infections (Figures 4 A-D). We tested if p38 MAPK could be an important player in SARS-CoV-2 replication by inhibiting p38 MAPK activity in SARS-CoV-2-infected Caco2 cells. Inhibition was confirmed by the reduced eIF4E phosphorylation (Figure 4E). Despite a substantial and significant drop in SARS-CoV-2 RNA in the supernatants of inhibitor-treated cultures (Figure 4F), the infectious titers remained unchanged (Figure 4 G). Unlike other reports (35,36), our results ruled out any significant impact of p38 on SARS-CoV-2 translation. Collectively, these results suggested that while p38 MAPK may be required for viral RNA replication, it does not play an active role in virus production.

**Figure 4:**
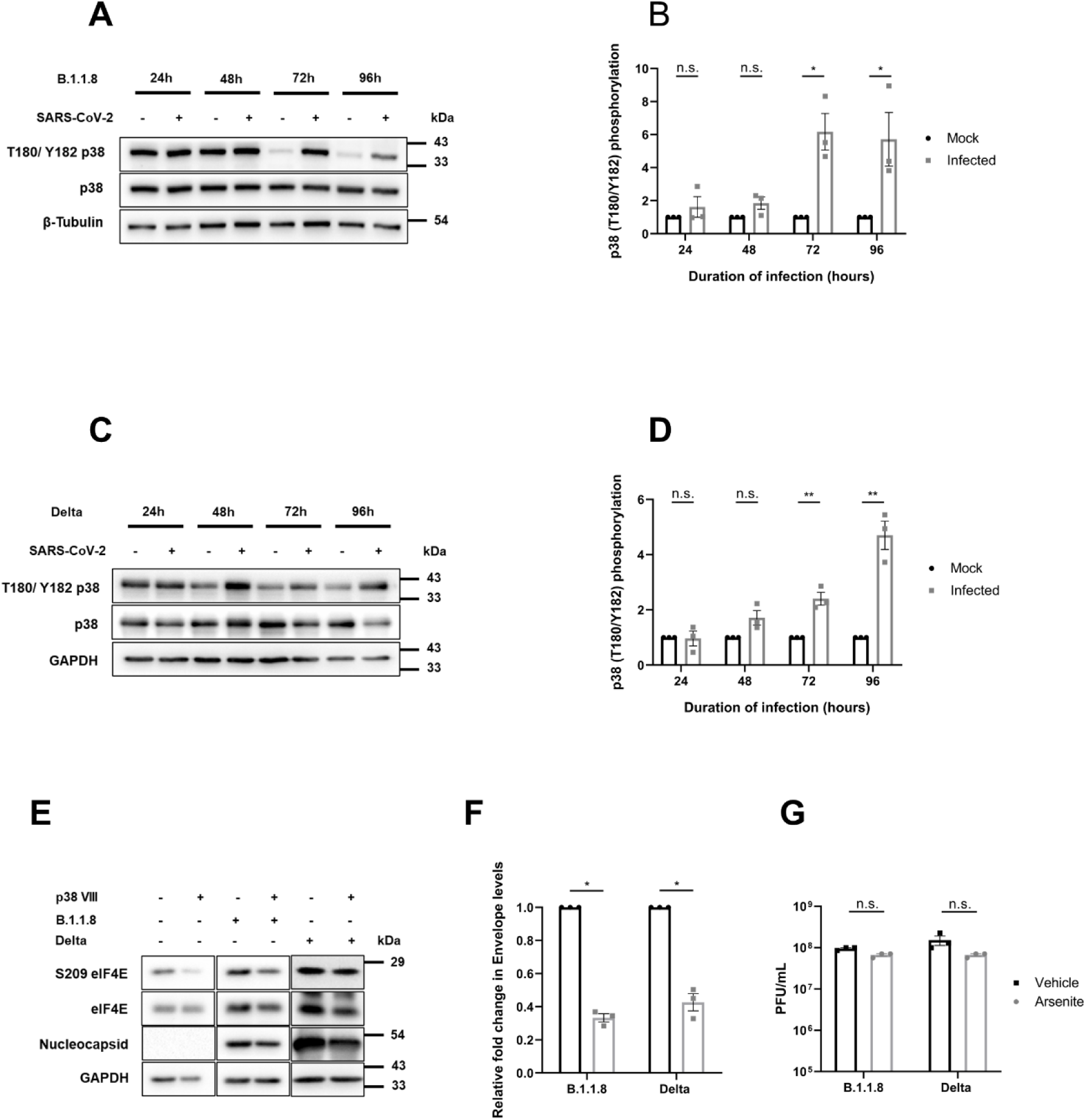
SARS-CoV-2 activates the p38 MAPK pathway independent of variants and this activation benefits viral replication. **(A, C)** Immunoblots showing changes in phosphorylation of p38-MAPK and its expression in Caco2 cells over different time points when infected with B.1.1.8 (A) and Delta (C), respectively. **(B, D)** Densitometric quantification of T180/Y182 phosphorylation from the panel in A (B) and panel in C (D), respectively. **(E)** SARS-CoV-2 variants infected Caco2 cells were treated with p38 VIII inhibitor (p38i) at 10 µM concentration for 24 hrs beginning at 24 hpi until harvesting. The inhibition was scored by the dephosphorylation status of eIF4E and viral protein abundance under p38 inhibited environment was also studied. **(F)** Relative fold change in SARS-CoV-2 extracellular E gene RNA, in DMSO and p38-inhibited cells. **(G)** Infectious viral titer measure in DMSO and p38-inhibited supernatants, quantified as PFU/mL. Graphs represent data from at least three sets and are plotted as mean ± SEM. p-values are represented as *, ** and ***, indicating p-values ≤ 0.05, 0.005 and 0.0005, respectively, n.s. represents non-significant.

### SARS-CoV-2 strongly inhibits mTORC1, but is unperturbed by the inhibition

As eIF2α and MAPK pathways were ruled out for their potential role in the polysome collapse during SARS-CoV-2 infection, we extended our studies to mTORC1. 4EBP1, a key substrate of mTORC1 and an important mediator of translation initiation, was used as the indicator of mTORC1 activity. The kinetics of 4EBP1 phosphorylation during SARS-CoV-2 infection in Caco2 cells was analyzed. We noticed a marked dephosphorylation of 4EBP1 in SARS-CoV-2 infected cells beyond 48 hpi (Figures 5 A-D), suggesting an inhibition of mTORC1 during infection. Both B.1.1.8 and Delta infections led to a comparable and sustained inhibition of mTORC1 activity, which coincided temporally with the observed polysome collapse. Since mTORC1 inhibition and polysome collapse shared similar time-frames, SARS-CoV-2 appears to inhibit mTORC1 in order to subvert the host translation machinery. The significance of this inhibition on the viral replication was further studied by inhibiting mTORC1 during the 24-hrs of infection by Rapamycin (Figure 5E). Though a noticeable drop in 4EBP1 phosphorylation was observed in Rapamycin treated cells, no significant change in N levels, or viral RNA or infectious titers were observed (Figures 5F and G). To further test the role of mTORC1 on SARS-CoV-2, we activated mTORC1 using MHY1485, and analyzed its impact on SARS-CoV-2 replication. Infected cells were treated with MHY1485 (10 μM) at 24 hpi and further continued up to 48 hpi. Despite a strong activation of mTORC1 and 4EBP1 phosphorylation with MHY1485, no significant change in viral RNA, N, or viral titers was observed (Figure 5H-J). These results confirmed that the inhibition of mTORC1 seen in SARS-CoV-2 infection has no consequence to the viral translation. Together, these results demonstrate that mTORC1 inhibition represents a conserved host response during infection and does not significantly alter viral replication efficiency.

**Figure 5:**
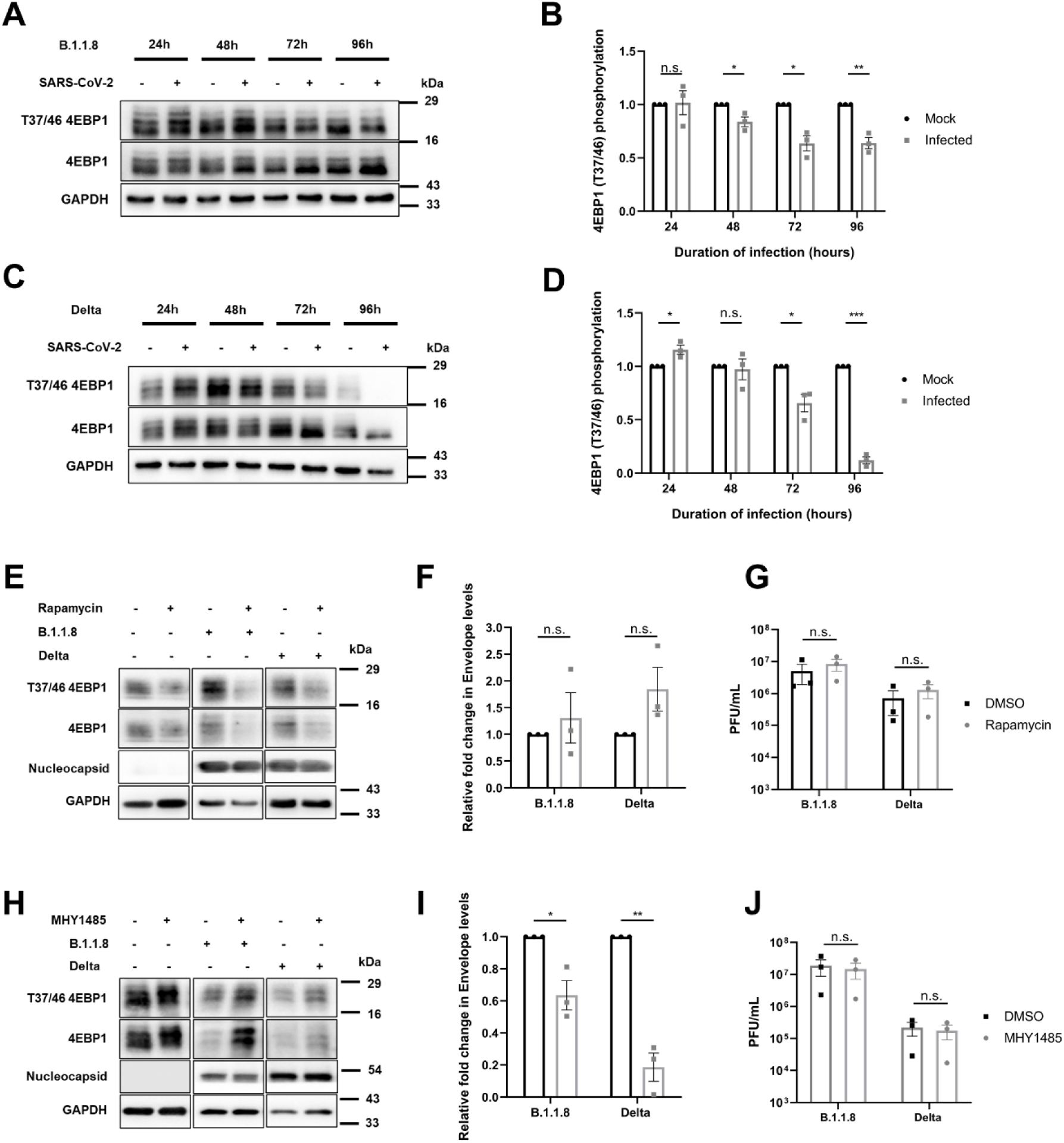
SARS-CoV-2 strongly inhibits mTORC1, but is unperturbed by the inhibition. **(A, C)** Immunoblots showing changes in phosphorylation of 4EBP1 and its expression in Caco2 cells over different time points when infected with B.1.1.8 (A) and Delta (C), respectively. **(B, D)** Densitometric quantification of T37/46 phosphorylation from the panel in A (B) and panel in C (D), respectively. **(E)** Caco2 cells were infected with SARS-CoV-2 variants for two hrs, treated with either DMSO or 25nM of Rapamycin up to 24 hpi, and harvested. A similar setup in mock cells was used. Inhibition was assessed by the drop in 4EBP1 phosphorylation in Rapamycin-treated mock and infected background. **(F)** Graph depicting relative fold changes in intracellular viral Envelope gene levels over rapamycin treatment in SARS-CoV-2 variant infected Caco2 cells. **(G)** Graph showing levels of viral titer with rapamycin treatment quantified as PFU/mL. **(H)** Mock and SARS-CoV-2 variants infected Caco2 cells were treated with either DMSO or 10 μM MHY1485 at 24 hpi and harvested at 48 hpi, and successful activation was assessed by increased phosphorylation of 4EBP1. **(I)** Graph showing relative fold changes in intracellular viral Envelope gene levels over MHY1485 treatment. **(J)** Graph showing levels of viral titer with MHY1485 treatment quantified as PFU/mL. Graphs represent data from at least three sets and are plotted as mean ± SEM. p-values are represented as *, ** and ***, indicating p-values ≤ 0.05, 0.005 and 0.0005, respectively, n.s. represents non-significant.

### 5’ TOP mRNAs escape translation repression and mTORC1 suppression

Polysome profiles of SARS-CoV-2 infected cultures showed no loss in overall ribosome abundance (Figures 1C and F). Translation of the mRNAs of ribosomal proteins (RPs) is regulated through the 5’ TOP element. Since the 5’ TOP mRNAs are regulated by mTORC1, we asked if the inhibition of mTORC1 pathway and translational repression negatively impacts ribosomal biogenesis by restricting the translation of the 5’ TOP transcripts.

We assessed cell lysates from mock- and SARS-CoV-2-infected cells for two ribosomal proteins, rpS3 (ribosomal protein S3) and rpL26 (ribosomal protein L26), as representatives for both subunits and 5’ TOP transcripts. Sustained levels of rpS3 were observed in the infected samples of both variants throughout the infection compared to the mock (B.1.1.8-Figures 6 A-B, Delta-Figures 6H-I). Notably, the levels of rpL26 increased with the progress of infection in the B.1.1.8 variant (B.1.1.8-Figures 6A-C, Delta-Figures 6H-J). Thus, despite a strong polysome dissociation and inhibition of mTORC1, ribosomal protein synthesis goes on unperturbed indicating that inhibition of mTORC1 activity is not affecting the translation of 5’ TOP mRNAs.

**Figure 6:**
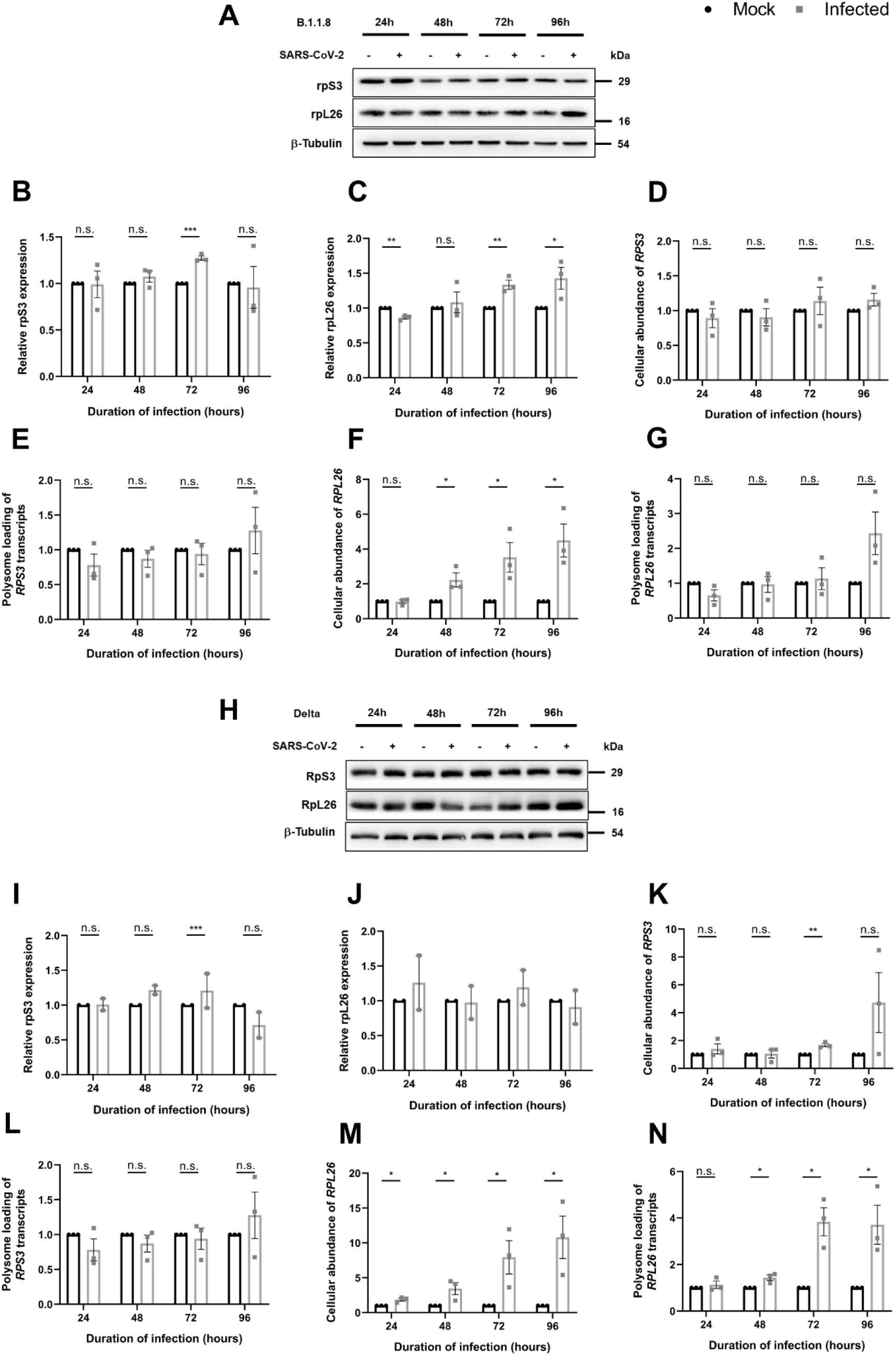
5’TOP mRNAs escape translation repression despite subdued mTORC1 activity during infection. **(A)** Immunoblot analysis of ribosomal protein rpS3 and rpL26 throughout infection with B.1.1.8. **(B, C)** Graphs showing densitometric quantification of rpS3 (B), and rpL26 (C) levels from the panel in A. **(D, E)** Cellular abundance of *RPS3* (D) and changes in its polysome association (E) with B.1.1.8 infection. **(F, G)** Cellular abundance of *RPL26* (F) and changes in its polysome association (G) with B.1.1.8 infection **(H)** Immunoblot showing the expression level of rpS3 and rpL26 under Delta infection in Caco2 cells **(I, J)** Graphs showing densitometric quantification of rpS3 and rpL26 levels from the panel in H, n=2. **(K, L)** Changes in cellular abundance of *RPS3* (K), and changes in its polysome association (L) with Delta infection**. (M, N)** Changes in cellular levels of *RPL26* (M) and changes in its polysome association with Delta infection. Graphs represent data from at least three sets until specified and are plotted as mean ± SEM. p-values are represented as *, ** and ***, indicating p-values ≤ 0.05, 0.005 and 0.0005, respectively, n.s. represents non-significant.

We next measured the cellular abundance and their ribosomal loading (association with translating ribosomes) of these two RP transcripts by quantitative RT-PCR. Cellular RNA and RNA from polysome fractions from across all time points after infection with both B.1.1.8 and Delta variants were analyzed. Steady-state levels of *RPS3* transcripts were observed in both B.1.1.8 and Delta variant infected cells (Figures 6D and K), and the transcripts were also associated with the polysomes throughout the infection (Figures 6E and L). Higher *RPL26* transcript levels were observed in the cells infected with both variants from 48 hpi (Figures 6 F and M). These transcripts were loaded onto polysome at a higher efficiency in cells infected with either of the variants, (Figures 6G and N). Subsequently, we studied the levels of viral RNA in cellular and polysome pools from B.1.1.8 and Delta infected Caco2 cultures in order to monitor the viral replication and the association of viral transcripts with the translating ribosomes. The abundance of viral RNA levels peaked at 48 hpi for both B.1.1.8 and Delta variants (Supplementary Figures S4A and C). A continuous association of viral RNAs with the polysomes was also observed in both variants (Supplementary Figures S4B and D), indicating unperturbed translation of viral RNA despite a significant depletion of the translating polysome pool. Overall, these results indicate that mTORC1 inhibition by SARS-CoV-2 infection does not affect the polysome loading of the viral RNA as well as ribosomal protein transcripts, and their translation, despite an mTORC1-suppressed environment.

## DISCUSSION

Several studies have demonstrated that SARS-CoV-2 infection suppresses host protein translation (3,5,7,14,37), with Nsp1 now being a well-documented driver of this effect by associating with 40S ribosomes to block host mRNA entry (7,16). Nsp2 also contributes to host immune suppression by stimulating viral translation even under hypoxic conditions, while specifically inhibiting interferon translation through interaction with specific translation initiation factors (38,39). Other viral proteins such as Orf9b significantly impact the innate immune system to regulate host responses, collectively inhibiting translation (7,34,40). However, the detailed picture of the complex, multilayered, regulatory networks that are frequently modulated by various viruses in order to establish their authority over the process of protein translation has not been established yet. Our study was aimed at understanding the overall picture of translation machinery and to gain insight into the signal pathways that normally regulate the machinery. We monitored polysome profiling to capture the polysome association in SARS-CoV-2 infected colon epithelial cell line Caco2 at multiple time intervals. These results were subsequently correlated with the modulation of the key regulatory pathways that control translation initiation. Using two variants of SARS-CoV-2, We identified that SARS-CoV-2 controls the translation machinery through the inhibition of mTORC1 pathway, while eIF2α and MAPK pathways are not affected in ways that could inhibit translation. Our results indicate that SARS-CoV-2 inhibits mTORC1, which is likely behind a massive collapse of polysomes thereby limiting translation of host transcripts. Interestingly, mTORC1 inhibition does not impact the polysome association of viral transcripts, and transcripts encoding ribosomal proteins. Our results find that the virus uses mechanisms to prevent polysome association that primarily targets host transcript while making sure that its own translation and those of ribosomal proteins are protected through mechanisms yet to be understood. The results were consistent between the two variants, highlighting the conserved mechanism targeted by SARS-CoV-2 in its pursuit to control host translation apparatus.

Similar to a previously published study (16), we did not observe any change in any of the three pathways that regulate translation during the early hours of infection, indicating any significant change on host translation signaling onsets much later. The only pathway that was impacted by SARS-CoV-2 was mTORC1. Unlike other reports that depict activated mTORC1 (41,42) we consistently observed an inhibition of this pathway with both the variants. Pharmacological activation of this pathway however did not have an impact on viral titers despite a curious drop in RNA levels, which indicates that the virus is able to sustain its translation despite an impact on its replication. Similar responses have been reported for other viruses such as Dengue (DENV). DENV translation, despite being cap-dependent, is unaffected by mTORC1 inhibition (43) Notwithstanding a strong inhibition of mTORC1 by the virus, translation of 5’ TOP transcripts, which are chiefly regulated by mTOR pathways, continued unhindered. This indicates a concerted effort by the virus to target specific transcripts regulated by mTORC1 without compromising on the requirements of the virus. It would be interesting to identify the host transcripts that are specifically targeted by the virus, which might lead to understanding the precise mechanisms by which this discrimination is established.

p38 and ERK1/2 MAPK signaling converge at cap-binding eIF4E to regulate translation initiation. Both kinases have a wide array of substrates that regulate a large number of processes including cell survival, stress, apoptosis, and interferon production. In our study, we noted that only ERK1/2, but not p38 MAPK, was dephosphorylated by SARS-CoV-2-mediated signaling activities. p38 activation remained sustained during both variant infections. The decreased viral replication upon p38 inhibition indicates SARS-CoV-2’s dependence on p38 activity for continued viral activities. This has been observed by other groups as well (44,45). Activated p38 promotes viral replication, influences innate immune signalling, and therefore viral infectivity (44). The impact of p38 inhibition on SARS-CoV-2 replication and inflammatory cytokine levels that has been observed by us and other groups (45) is possibly why its activation status is unaffected throughout evolution of the virus. Between the activated p38 MAPK and dephosphorylated state of ERK1/2, the lack of significant impact on eIF4E phosphorylation is expected. Since eIF4E phosphorylation is understood to affect only a select set of mRNAs translationally (46), we believe that its contribution to the global suppression of translation activities caused by SARS-CoV-2 infection could be limited and more studies are necessary to determine its impact.

It is fascinating to note that SARS-CoV-2 infection manages to avoid eIF2α phosphorylation. eIF2α is phosphorylated by one of its four kinases most of which are activated upon various stress exerted on the cell. RNA viruses often impart intense stress on ER that is relayed to PERK (47,48). PKR, one of the dsRNA sensors is often activated by RNA viral infections. These observations indicate that SARS-CoV-2 depends on the canonical mechanism of translation initiation that requires the availability of active ternary complexes, which eIF2α is a part of. Intriguingly, one report found that PKR inhibition could work as a potent antiviral against SARS-CoV-2(49), suggesting that despite any purported effect on eIF2α phosphorylation in our study, its kinases may still be involved in curbing SARS-CoV-2 infection. Since eIF2α phosphorylation results in the inhibition of new initiation events that would adversely affect the translation of viral transcripts as well, SARS-CoV-2 might have evolved strategies to bypass this modification. One such report suggests that Nsp3 and Nsp4 of SARS-CoV-2 suppress PERK, which is also involved in the unfolded protein response pathway (50), which could also impact global translation through eIF2α. Another report indicates that SARS-CoV-2 inhibits stress granule assembly, a process largely regulated by eIF2α, and this may be a mechanism to allow continued viral replication (51). Together, we believe that the inhibited status of eIF2α found in both variants during active infection allows viral translation to continue unimpeded.

To summarize our findings, we observe the complex relation between host translation regulation pathways in response to viral infection. The most striking impact was observed on mTOR pathway although this did not deter translation of certain key mTOR substrates such as 5’TOP transcripts. Surprisingly, unlike other RNA viruses, eIF2α phosphorylation does not seem to be sustained or increased along the course of infection. Our study focuses on the interplay between host and viral systems after the establishment of active infection, as opposed to other studies that have focused on its immediate impact. Our observations clarify that there is a dynamic rewiring in cellular signalling in response to virus-mediated effects on translation. Findings from our lab also indicate at novel means by which SARS-CoV-2 evades host innate immune responses as well that may also explain how some of our observations differ from other RNA viruses (Tandel et al).

## LIMITATIONS

While this study establishes a conserved mechanism of host translational suppression by distinct SARS-CoV-2 variants, it is limited by the use of a single epithelial cell model, which may not fully capture the heterogeneity of infection across different tissue environments. The experiments were performed with pre-Omicron lineages, and it remains to be determined whether newer variants exhibiting altered pathogenicity retain similar translational control or not. Furthermore, although our findings clearly links mTORC1 inhibition with polysome collapse, the specific viral or host determinants driving this effect are not dissected here and needs further investigation. Nonetheless, by integrating polysome profiling with signalling analyses, this study provides a comprehensive framework that advances understanding of how SARS-CoV-2 modulates host translation.

### Experimental procedures Antibodies and inhibitors

All primary antibodies were purchased from Cell Signaling Technologies except the GAPDH and anti-SARS-CoV-2 Nucleocapsid were procured from Thermo Fisher Scientific. HRP-conjugated anti-rabbit and anti-mouse secondary antibodies were purchased from Jackson ImmunoResearch. Rapamycin and MHY1485 were from Sigma, whereas the p38 VIII inhibitor and Sodium arsenite were from Merck Millipore.

### Cell culture

Caco2, human colorectal adenocarcinoma epithelial cells were procured from ATCC and were cultured in Dulbecco’s Modified Eagle’s Medium (DMEM; from Gibco) with 20% Fetal Bovine Serum (FBS; Hyclone), 1× penicillin-streptomycin cocktail (Pen-strep, Gibco), and 1× non-essential amino acids (NEAA, Gibco), at 37°C and 5% CO_2_. Cells were continuously passaged at 70-80% confluency and mycoplasma contamination was monitored periodically.

### SARS-CoV-2 Infection and quantification

Two Indian isolates of SARS-CoV-2 strains were used in this study (GSSAID id: EPI_ISL_458046, (B.1.1.8), and EPI_ISL_2775201 (Delta) (52,53). All the viral cultures were propagated in Vero (CCL-81) cells in serum and antibiotics-free conditions. Caco2 cells were infected at 1 MOI for 3 hrs in serum-free conditions after which the media was replaced with complete media and further incubated until the time of harvesting as per experiment requirement. At the time of harvesting, the cells were first trypsinized and collected separately for protein and RNA study. The extracellular RNA from the supernatants was isolated (MACHEREY-NAGEL GmbH & Co. KG) and the SARS-CoV-2 RNA was quantified using a commercial kit (LabGun™ COVID-19 RT-PCR Kit) following manufacturers protocol in Roche LightCycler 480. The viral E gene was normalized with internal control provided in the kit, and fold changes were calculated using 2^(-ΔΔCt)^ method. The infectious viral particle numbers in the cell culture supernatant were quantified using the plaque-forming unit (PFU/mL) assay. Briefly, the supernatant was log diluted (10^-1^–10^-7^) in 1× serum-free DMEM and used for infecting Vero monolayer-grown twelve-well plates. 3 hpi, the cells were briefly washed and were overlaid with agarose: DMEM mix (in 1:1 ratio; 2 x DMEM with 5% FBS and 1% penicillin-streptomycin mixed with equal volumes of 2% LMA), after which the plates were incubated undisturbed for 6 days at 37°C. Later, the cells were fixed with 4% formaldehyde and stained with crystal violet. The clear zones were counted and PFU was calculated as PFU/mL.

### Inhibitions and infection

For the inhibition experiments, 0.45 × 10^6^ Caco2 cells were seeded in a six-well format and 24 hrs later the cells were infected with SARS-CoV-2 variants, at 1 MOI for 3 hrs in serum-free media. For rapamycin inhibition, the infection media was replaced with serum-sufficient media containing 25 nM Rapamycin or DMSO, and incubated for 22 hrs. At the end of the treatment, the cells were harvested and protein and RNA were prepared. For the activation of mTORC1, cells were treated with 10 µM MHY1485 or DMSO at 24 hpi and harvested at 48 hpi. The p38 inhibition was carried out in a similar fashion to the MHY1485 experiment, at a final concentration of 10 µM. For ISR activation, SARS-CoV-2 variant infected cells were treated with 10 µM of sodium arsenite or water for 8 hrs at 16 hpi. The intracellular and extracellular RNA were subjected to qRT-PCR, and the protein lysates were subjected to western blotting for confirmation of inhibition or activation.

### Polysome preparation and RNA precipitation

Polysomes were fractionated as explained elsewhere (54). Caco2 cells were grown in 175cm^2^ flasks till 70% confluency and subsequently infected with SARS-CoV-2 variants at 1 MOI. Media was changed after 2 hrs, and cells were harvested at 24, 48, 72, and 96 hpi, along with mock-infected cells grown alongside for each time point. The cells were incubated for 5-10 minutes, harvested, and washed twice with a solution of ice-cold 1×PBS containing 100 μg/mL cycloheximide, to freeze the polysomes on the mRNAs. They were subsequently lysed in polysome lysis buffer containing 20 mM Tris-Cl pH 8.0, 140 mM KCl, 1.5 mM MgCl_2_, 0.5 mM DTT, 1% Triton X-100, 1× protease inhibitor, 0.5 mg/L heparin,100 μg/mL cycloheximide, and RNase inhibitor. Crude RNA was quantified using a spectrophotometer, and 90 μg was layered onto 11 mL of 10-50% linear sucrose gradient (20 mM Tris-Cl pH 8.0, 140 mM KCl, 1.5 mM MgCl_2_, 0.5 mM DTT, 100 μg/mL cycloheximide, 1mM PMSF, 10-50% sucrose). The resulting gradients were centrifuged in an SW41 Ti rotor (Beckman Coulter) at 35,000 r.p.m. at 4°C for 3.5 hrs. The polysome samples were fractionated using the Teledyne ISCO fraction collector system and absorbance was measured and graphically noted at 254 nM. Polysome profiles of mock and infected cells for each time point were digitized and overlaid using Inkscape. Later, total polysome RNA was precipitated by adding 1/10^th^ volume of 3M sodium acetate and an equal volume of absolute ethanol, incubated at -20°C and centrifuged at 16000g, and the pellet obtained was airdried and resuspended in DEPC water. To this, 4M LiCl was added to a final concentration of 2.5M, incubated at -20°C overnight. After centrifugation, the pellet was washed twice with 70% ethanol, airdried and resuspended in DEPC water. Subsequently, the RNA samples were purified using MN NucleoSpin RNA kit following manufacturer’s protocol. The abundance of polysome-associated mRNAs were quantified through qRT-PCR.

### Quantitative RT-PCR

Cellular and polysomal RNA were extracted using the Nucleospin RNA kit (MACHEREY-NAGEL) according to the manufacturer’s instructions. Equal quantities of RNA were reverse-transcribed into cDNA using the Primescript Reverse transcriptase kit (Takara) as per the manufacturer’s protocol. Subsequently, equal amounts of cDNA were used for specific gene expression quantification using specific primers (Table 1) and the SYBR Green Master Mix (Takara) on a LightCycler 480 system (Roche). Host-specific transcripts were normalized to GAPDH, while viral transcripts were normalized using the internal control gene included in the LabGun™ COVID-19 RT-PCR Kit. Relative fold changes between control and experimental samples were calculated using the 2^(-ΔΔCt)^ method and presented as graphs.

**Table 1.**
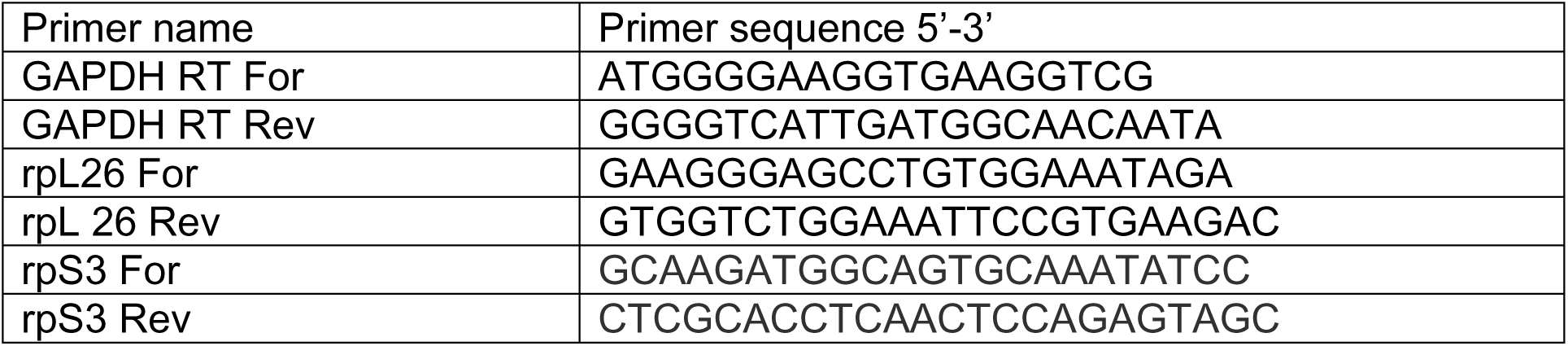

### Immunoblotting

Protein pellets were lysed in 1 × Nonidet P-40 lysis buffer (1% Nonidet P-40, 50 mM Tris-HCl, 150 mM NaCl (pH 7.5), EGTA, 1 mM sodium orthovanadate, 10 mM sodium pyrophosphate, 100 mM NaF, and 1 mM PMSF) incubated on ice for 20 minutes with intermittent vortexing and centrifuged at 13,000 rpm for 15 minutes at 4°C. The supernatants containing the proteins were collected and quantified using BCA reagents (G Biosciences). Lysates were mixed with 6× denaturing dye and the proteins were resolved using SDS-PAGE and transferred to PVDF membranes. The membranes were blocked in 5% BSA dissolved in 1× TBST before the addition of primary antibodies. Primary antibodies against the proteins of interest were diluted in the blocking buffer, added to the membrane, and incubated overnight at 4°C. Later, the membranes were washed in 1× TBST, secondary antibodies conjugated with HRP were added and the blots were developed on a Bio-Rad Chemidoc MP system using SuperSignal West Pico PLUS (Thermo Fisher) and SuperSignal West Femto Maximum Sensitivity (Thermo Fisher) chemiluminescent substrate kits.

### Statistical analysis

For each experiment, at least three independent replicates (until specified) were used to calculate mean ± SEM and plotted graphically wherever indicated. Statistical significance was measured using a two-tailed, unpaired Student t-test, and the resultant p values were represented as *, **, *** indicating p values ≤ 0.05, 0.005, and 0.0005, respectively.

## Data availability

All data pertaining to this manuscript are included within the manuscript.

## Institutional biosafety

Institutional biosafety clearance was obtained by K.H.H., for the experiments pertaining to SARS-CoV-2.

## Supporting information

This article contains supporting information.

## Acknowledgements

We thank Mohan Singh Moodu and Amit Kumar for assisting with logistics and Karthika S Nair for her assistance with some experiments.

## Contributions

The experiments were conceived by H.P., D.G., and K.H.H. H.P., and D.G., performed polysome profiling, and qRT-PCRs. D.G., D.K., and V.S prepared infectious SARS-CoV-2, performed infections, quantified them and analyzed data. H.P., A.P.S., D.G., D.K., and A.H.S. performed immunoblotting. H.P., and D.G., assisted K.H.H in writing the manuscript.

