## Supplemental Figures 1-4 for "SARS-CoV-2 Infection Causes Strong mTORC1 Inhibition and Massive Polysome Collapse"

#### Figure S1

**A**

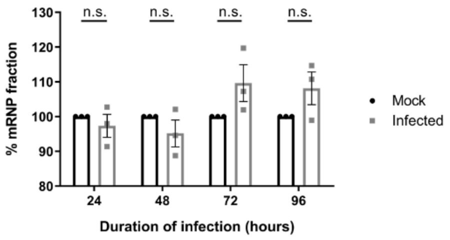


**B**

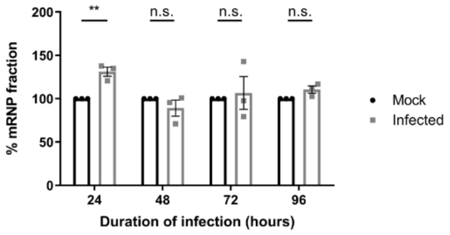


**Figure S1: B.1.1.8 and Delta infections do not impact the cellular polysome-free mRNP fractions (A)** and **(B)** Graphs showing changes in mRNP fraction over 24, 48, 72 and 96 hpi with B.1.1.8 and Delta infection, respectively. The area under the curve was calculated using ImageJ software. Area corresponding to mRNPs was normalized to the profile's total area. Graphs represent data from three sets and are plotted as mean ± SEM. p-values are represented as n.s (not significant), and ** indicating p-values ≤ 0.005.

### Figure S2

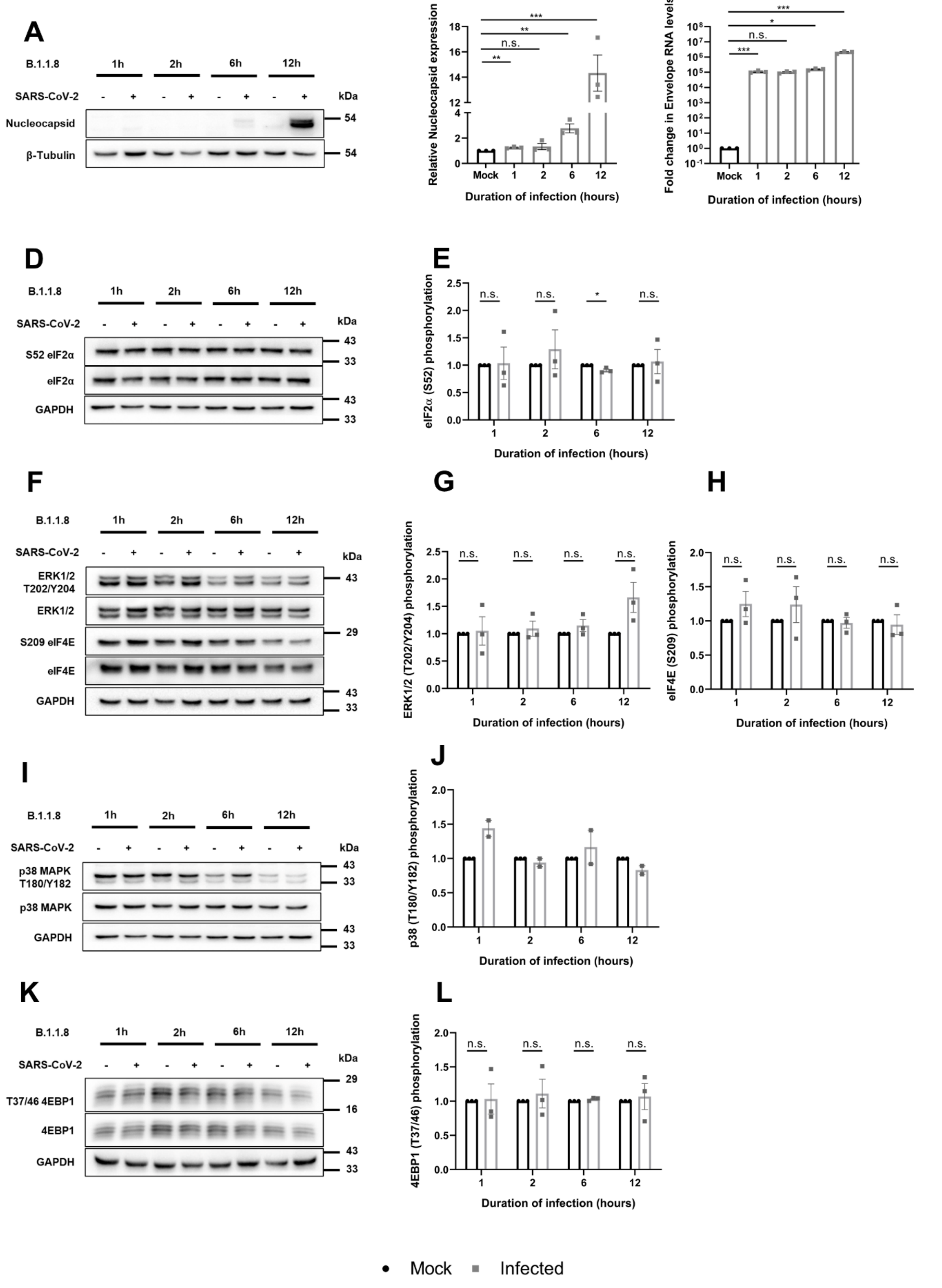
A
B.1.1.8
1h
2h
6h
12h
SARS-CoV-2
kDa
Nucleocapsid
β-Tubulin
54
B
Relative Nucleocapsid expression
Duration of infection (hours)
Mock
C
Fold change in Envelope RNA levels
D
S52 eIF2α
eIF2α
GAPDH
E
eIF2α (S52) phosphorylation
F
ERK1/2 T202/Y204
ERK1/2
S209 eIF4E
eIF4E
G
ERK1/2 (T202/Y204) phosphorylation
H
eIF4E (S209) phosphorylation
I
p38 MAPK T180/Y182
p38 MAPK
J
p38 (T180/Y182) phosphorylation
K
T37/46 4EBP1
4EBP1
L
4EBP1 (T37/46) phosphorylation
n.s.
Mock
Infected

**Figure S2: B.1.1.8 does not affect translation regulatory pathways during the early hours of infection. (A)** Immunoblot analysis of Caco2 infected cells with 1 MOI of B.1.1.8 for 1, 2, 6, and 12 hrs along with mock-infected controls, showing levels of nucleocapsid protein. **(B)** Densitometry of Nucleocapsid expression normalized to β-Tubulin. **(C)** Graph depicting relative fold changes in SARS-CoV-2 E gene, quantified through real-time PCR, across early hours of infection with B.1.1.8. **(D)** Immunoblot showing the kinetics of eIF2α phosphorylation at S52 and its expression levels. **(E)** Densitometry of eIF2α phosphorylation at S52. **(F)** Immunoblot showing phosphorylation of ERK1/2 at T202/Y204 and eIF4E at S209, and their expression. **(G)** and **(H)** Densitometric analysis of ERK1/2 and eIF4E phosphorylation, respectively. **(I)** Immunoblot showing phosphorylation of p38 at T180/Y182 and its expression levels. **(J)** Densitometry of p38 phosphorylation across early time points, n=2. **(K)** Immunoblot showing 4EBP1 phosphorylation at T37/46 and its expression across four time points. **(L)** Densitometry of 4EBP1 phosphorylation. All densitometric analyses showing changes in phosphorylated proteins are normalized with their respective total proteins and GAPDH. The infected values were then normalized to the respective mock values. All the graphs represent data from at least three sets, unless specified, and are plotted as mean ± SEM. p-values are represented as n.s (not significant), *, ** and ***, indicating p-values ≤ 0.05, 0.005, and 0.0005, respectively.

### Figure S3

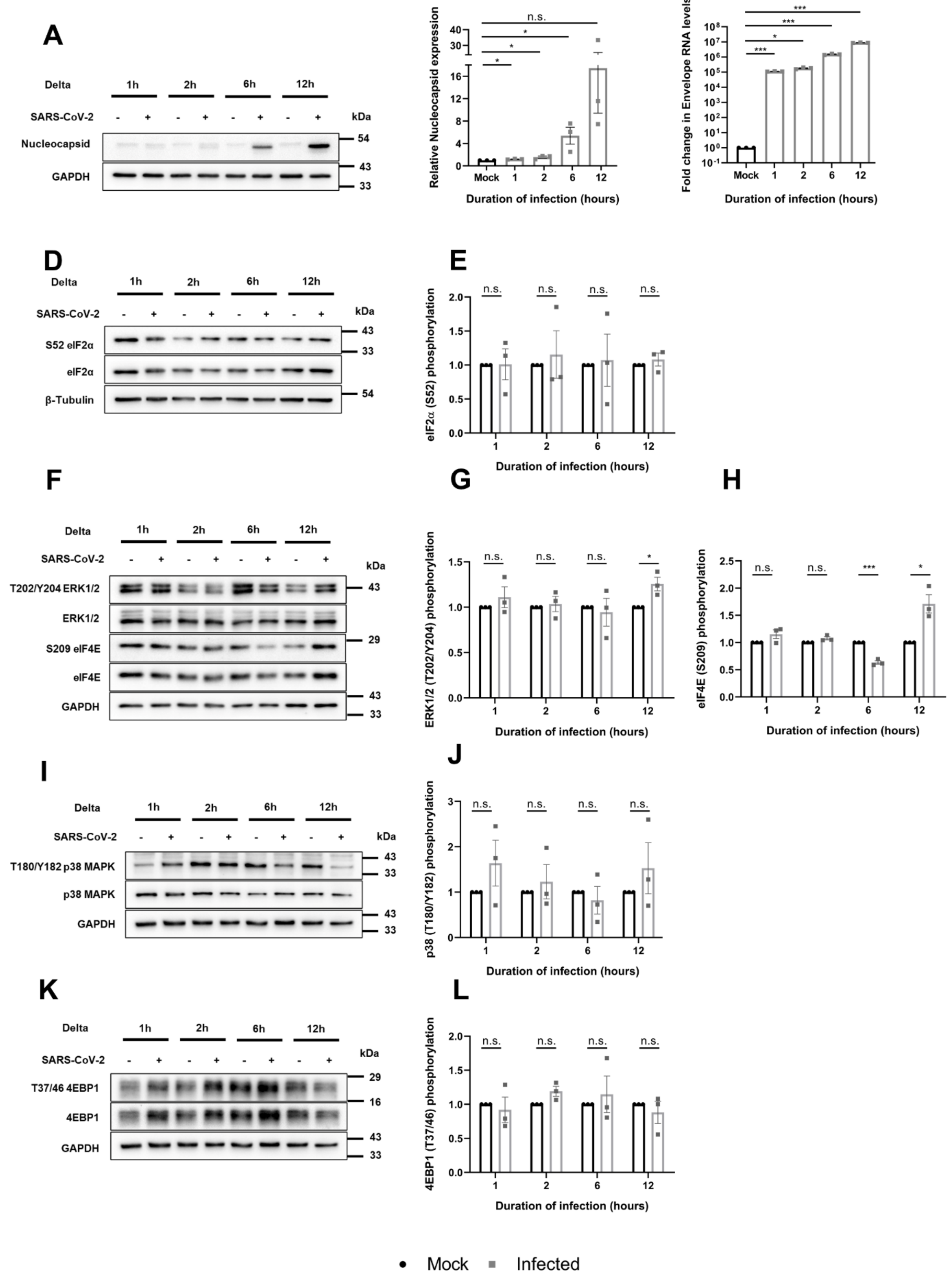
A
Delta
1h
2h
6h
12h
SARS-CoV-2
kDa
Nucleocapsid
GAPDH
54
43
33
B
Relative Nucleocapsid expression
Duration of infection (hours)
Mock
n.s.
C
Fold change in Envelope RNA levels
Duration of infection (hours)
D
S52 eIF2α
eIF2α
β-Tubulin
E
eIF2α (S52) phosphorylation
F
T202/Y204 ERK1/2
ERK1/2
S209 eIF4E
eIF4E
G
ERK1/2 (T202/Y204) phosphorylation
H
eIF4E (S209) phosphorylation
I
T180/Y182 p38 MAPK
p38 MAPK
J
p38 (T180/Y182) phosphorylation
K
T37/46 4EBP1
4EBP1
L
4EBP1 (T37/46) phosphorylation
Mock
Infected

**Figure S3: Delta does not affect translation regulatory pathways during the early hours of infection. (A)** Immunoblot analysis of Caco2 infected cells with 1 MOI of Delta for 1, 2, 6, and 12 hrs along with mock-infected controls, showing levels of nucleocapsid protein. **(B)** Densitometry of Nucleocapsid expression normalized to β-Tubulin. **(C)** Graph depicting relative fold changes in SARS-CoV-2 E gene, quantified through real-time PCR, across early hours of infection with Delta. **(D)** Immunoblot showing the kinetics of eIF2α phosphorylation at S52 and its expression levels. **(E)** Densitometry of eIF2α phosphorylation at S52. **(F)** Immunoblot showing phosphorylation of ERK1/2 at T202/Y204 and eIF4E at S209, and their expression. **(G, H)** Densitometric analysis of ERK1/2 (G) and eIF4E phosphorylation (H), respectively. **(I)** Immunoblot showing phosphorylation of p38 at T180/Y182 and its expression levels. **(J)** Densitometry of p38 phosphorylation across early time points. **(K)** Immunoblot showing 4EBP1 phosphorylation at T37/46 and its expression across four time points. **(L)** Densitometry of 4EBP1 phosphorylation. All densitometric analyses showing changes in phosphorylated proteins are normalized with their respective total proteins and GAPDH, and infected values were then normalized to their respective mock values. All the graphs represent data from at least three sets, and are plotted as mean ± SEM. p-values are represented as * and ***, indicating p-values ≤ 0.05 and 0.0005, respectively.

### Figure S4

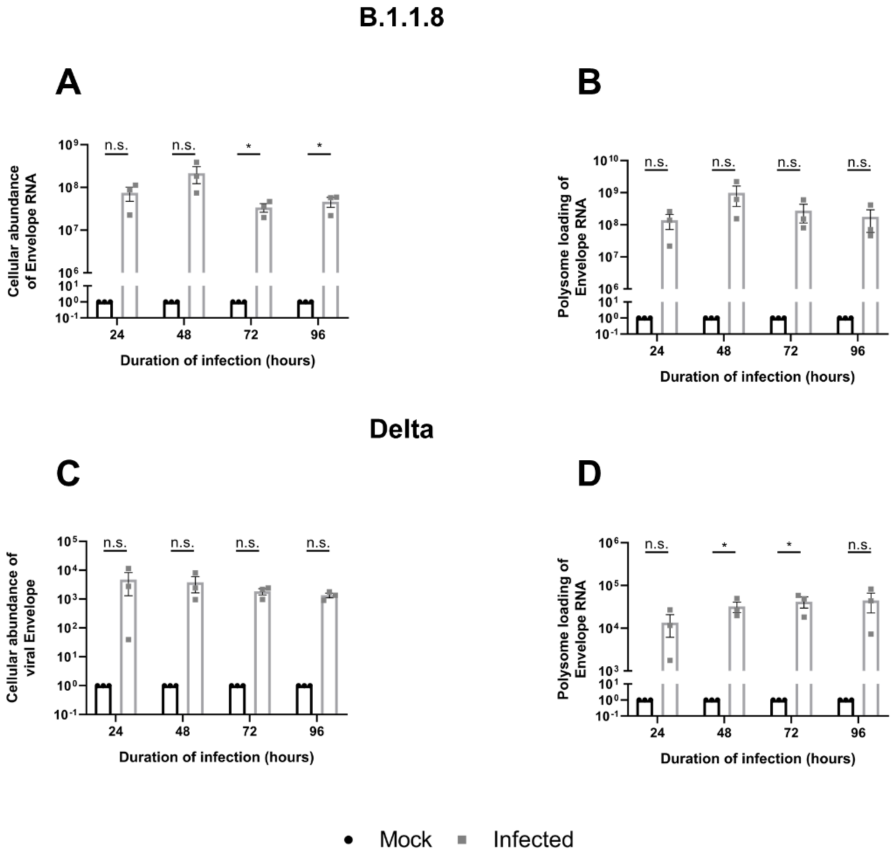


**Figure S4: SARS-CoV-2 viral RNA are enriched in cellular and polysome fractions. (A)** Graph showing changes in SARS-CoV-2 Envelope levels in the cellular pool with B.1.1.8 infection. **(B)** Graph depicting SARS-CoV-2 Envelope levels in polysome fraction with B.1.1.8 infection. **(C)** Graph representing the cellular abundance of the viral Envelope gene over different time intervals with Delta infection. **(D)** Graph showing polysome loading of Envelope RNA on Delta infection. Graphs represent data from at least three sets and are plotted as mean ± SEM. p-values are represented as * indicating p-values ≤ 0.05, n.s. represents non-significant.
